# Maternal and fetal genetic contribution to gestational weight gain

**DOI:** 10.1101/116434

**Authors:** Nicole M Warrington, Rebecca Richmond, Bjarke Fenstra, Ronny Myhre, Romy Gaillard, Lavinia Paternoster, Carol A Wang, Robin N Beaumont, Shikta Das, Mario Murcia, Sheila J Barton, Ana Espinosa, Elisabeth Theiring, Mustafa Atalay, Niina Pitkänen, Ioanna Ntalla, Anna E Jonsson, Rachel Freathy, Ville Karhunen, Carla MT Tiesler, Catherine Allard, Andrew Crawford, Susan M Ring, Mads Melbye, Per Magnus, Fernando Rivadeneira, Line Skotte, Torben Hansen, Julie Marsh, Mònica Guxens, John W Holloway, Harald Grallert, Vincent WV Jaddoe, William L Lowe, Theano Roumeliotaki, Andrew T Hattersley, Virpi Lindi, Katja Pahkala, Kalliope Panoutsopoulou, Marie Standl, Claudia Flexeder, Luigi Bouchard, Ellen Aagard Nohr, Loreto Santa Marina, Manolis Kogevinas, Harri Niinikoski, George Dedoussis, Joachim H Heinrich, Rebecca M Reynolds, Timo Lakka, Eleftheria Zeggini, Olli T Raitakari, Leda Chatzi, Hazel M Inskip, Mariona Bustamante Pineda, Marie-France Hivert, Marjo-Riitta Jarvelin, Thorkild IA Sørensen, Craig Pennell, Janine F Felix, Bo Jacobsson, Frank Geller, David M Evans, Debbie A Lawlor, Early Growth Genetics (EGG) consortium

**Affiliations:** University of Queensland Diamantina Institute, Translational Research Institute, Brisbane, Australia; School of Women’s and Infants’ Health, The University of Western Australia, Perth, Australia; Medical Research Council Integrative Epidemiology Unit at the University of Bristol, Bristol, UK; School of Social and Community Medicine, University of Bristol, Bristol; Department of Epidemiology Research, Statens Serum Institute, Copenhagen, Denmark; Norwegian Institute of Public Health, Oslo, Norway; The Generation R Study Group, Erasmus MC, University Medical Center Rotterdam, Rotterdam, the Netherlands; Department of Epidemiology, Erasmus MC, University Medical Center Rotterdam, Rotterdam, the Netherlands; Department of Pediatrics, Erasmus MC, University Medical Center Rotterdam, Rotterdam, the Netherlands; Institute of Biomedical and Clinical Science, University of Exeter Medical School, Royal Devon and Exeter Hospital, Exeter, UK; Department of Public Health and Primary Care, School of Public Health, Imperial College London, London, UK.; Epidemiology and Environmental Health Joint Research Unit, FISABIO–Universitat Jaume I–Universitat de València, Valencia, Spain; Spanish Consortium for Research on Epidemiology and Public Health (CIBERESP), Spain; MRC Lifecourse Epidemiology Unit, Faulty of Medicine, University of Southampton, Southampton, UK; ISGlobal, Centre for Research in Environmental Epidemiology (CREAL), Barcelona, Spain; IMIM (Hospital del Mar Medical Research Institute), Barcelona, Spain; Universitat Pompeu Fabra (UPF), Barcelona, Spain; Institute of Epidemiology I, Helmholtz Zentrum München- German Research Center for Environmental Health, Neuherberg, Germany; Division of Metabolic and Nutritional Medicine, Dr. von Hauner Children’s Hospital, University of Munich Medical Center, Munich, Germany; Institute of Biomedicine, School of Medicine, University of Eastern Finland, Kuopio, Finland; Research Centre of Applied and Preventive Cardiovascular Medicine, University of Turku, Turku, Finland; William Harvey Research Institute, Barts and the London School of Medicine and Dentistry, Queen Mary University of London, London, UK; The Novo Nordisk Foundation Center for Basic Metabolic Research, Section of Metabolic Genetics, Faculty of Health and Medical Sciences, University of Copenhagen, Copenhagen, Denmark; Center for Life Course Health Research, Faculty of Medicine, University of Oulu, Oulu, Finland.; Biocenter Oulu, University of Oulu, Finland.; Centre de Recherche du Centre Hospitalier Universitaire de Sherbrooke, Sherbrooke, Canada; British Heart Foundation Centre for Cardiovascular Science, Queen’s Medical Research Institute, University of Edinburgh, Edinburgh, UK; ALSPAC (Children of the 90s), School of Social and Community Medicine, University of Bristol, Bristol; Department of Clinical Medicine, Copenhagen University, Copenhagen, Denmark; Department of Medicine, Stanford School of Medicine, Stanford, California, USA; Department of Internal Medicine, Erasmus MC, University Medical Center Rotterdam, Rotterdam, the Netherlands; Department of Child and Adolescent Psychiatry/Psychology, Erasmus University Medical Centre–Sophia Children’s Hospital, Rotterdam, the Netherlands; Human Development and Health, Faculty of Medicine, University of Southampton, Southampton, UK; Institute of Epidemiology II, Research Unit of Molecular Epidemiology, Helmholtz Zentrum München Research Center for Environmental Health, Neuherberg, Germany; German Center for Diabetes Research (DZD), Neuherberg, Germany; Clinical Cooperation Group Type 2 Diabetes, Helmholtz Zentrum München, Neuherberg, Germany; Clinical Cooperation Group Nutrigenomics and Type 2 Diabetes, Helmholtz Zentrum München, Neuherberg, Germany and Technische Universität München, Freising, Germany; Department of Medicine, Division of Endocrinology, Metabolism, and Molecular Medicine, Feinberg School of Medicine, Northwestern University, Chicago, USA; Department of Social Medicine, University of Crete, Greece; Paavo Nurmi Centre, Sports and Exercise Medicine Unit, Department of Health and Physical Activity, Turku, Finland; Department of Human Genetics, Wellcome Trust Sanger Institute, Hinxton, UK; Department of Biochemistry, Faculty of medicine and life sciences, Université de Sherbrooke, Sherbrooke, Canada; Research Unit for Gynaecology and Obstetrics, Department of Clinical Research, University of Southern Denmark, Denmark; Public Health Division of Gipuzkoa, Basque Government, Spain; Health Research Institute, Biodonostia, San Sebastián, Spain; Department of Pediatrics, Turku University Hospital, Turku, Finland; Department of Nutrition & Dietetics, Harokopio University of Athens, Athens, Greece; Institute and Outpatient Clinic for Occupational, Social and Environmental Medicine, Inner City Clinic, University Hospital Munich, Ludwig Maximilian University of Munich, Munich, Germany; Department of Clinical Physiology and Nuclear Medicine, Kuopio University Hospital, School of Medicine, University of Eastern Finland, Kuopio, Finland; Kuopio Research Institute of Exercise Medicine, Kuopio, Finland; Department of Clinical Physiology and Nuclear Medicine, Turku University Hospital, Turku, Finland; NIHR Southampton Biomedical Research Centre, University of Southampton and University Hospital Southampton NHS Foundation Trust, Southampton, UK; Centre for Genomic Regulation (CRG), The Barcelona Institute of Science and Technology, Barcelona, Spain; Department of Population Medicine at Harvard Pilgrim Health Care Institute, Harvard Medical School, Boston, USA; Diabetes Unit, Massachusetts General Hospital, Boston, USA; Department of Epidemiology and Biostatistics, MRC–PHE Centre for Environment & Health, School of Public Health, Imperial College London, London, UK.; Unit of Primary Care, Oulu University Hospital, Oulu; Institute of Preventive Medicine, Bispebjerg and Frederiksberg Hospital, The Capital Region, Copenhagen, Denmark; Department of Obstetrics and Gynecology, Sahlgrenska Academy, Gothenburg University, Gothenburg, Sweden; Department of Genetics and Bioinformatics, Domain of Health Data and Digitalization, Institute of Public Health, Oslo, Norway

## Abstract

**Background:** Clinical recommendations to limit gestational weight gain (GWG) imply high GWG is causally related to adverse outcomes in mother or offspring, but GWG is the sum of several inter-related complex phenotypes (maternal fat deposition and vascular expansion, placenta, amniotic fluid and fetal growth). Understanding the genetic contribution to GWG could help clarify the potential effect of its different components on maternal and offspring health. Here we explore the genetic contribution to total, early and late GWG.

**Participants and Methods:** A genome-wide association study was used to identify maternal and fetal variants contributing to GWG in up to 10,543 mothers and up to 16,317 offspring of European origin, with replication in 10,660 mothers and 7,561 offspring. Additional analyses determined the proportion of variability in GWG from maternal and fetal common genetic variants and the overlap of established genome-wide significant variants for phenotypes relevant to GWG (e.g. maternal BMI and glucose, birthweight).

**Results:** We found that approximately 20% of the variability in GWG was tagged by common maternal genetic variants, and that the fetal genome made a surprisingly minor contribution to explaining variation in GWG. We were unable to identify any genetic variants that reached genome-wide levels of significance (P<5×10^−8^) and replicated. Some established maternal variants associated with increased BMI, fasting glucose and type 2 diabetes were associated with lower early, and higher later GWG. Maternal variants related to higher systolic blood pressure were related to lower late GWG. Established maternal and fetal birthweight variants were largely unrelated to GWG.

**Conclusion:** We found a modest contribution of maternal common variants to GWG and some overlap of maternal BMI, glucose and type 2 diabetes variants with GWG. These findings suggest that associations between GWG and later offspring/maternal outcomes may be due to the relationship of maternal BMI and diabetes with GWG.

## Introduction

High and low levels of gestational weight gain (GWG), defined as the weight a woman gains during pregnancy before delivery of her infant,^1^ are associated with a wide range of adverse outcomes for mother and child in the short- (during pregnancy and the perinatal period), and longer-term.^2-9^ As a consequence of these associations, recommendations for healthy GWG are increasingly used in antenatal care,^1^ despite a lack of evidence that any of these associations are causal, and if they are, what the mechanisms underlying them might be.^3,10^ GWG is a complex phenotype that is influenced by maternal responses to pregnancy, such as gestational fat deposition and volume expansion, as well as fetal growth, placental size and amniotic fluid volume.^3,10^ Each of these are likely to be influenced both by maternal and fetal genes and environmental exposures. Understanding the maternal and fetal genetic contributions to GWG could shed light on both genetic and non-genetic contributions to between woman variation in GWG.^11,12^ For example, we have recently used maternal genetic instrumental variables to determine the causal effect of maternal pregnancy adiposity and related traits on offspring birthweight and ponderal index.^13^

Amongst 1,159 European origin Swedish maternal twin pairs (694 pairs with data on their first pregnancies and 465 on their second) it has been shown that approximately 40% of variability in first pregnancy GWG was due to genetic factors.^14^ Other studies have examined the associations of candidate maternal and/or fetal adiposity or diabetes related genetic variants with GWG and yielded inconsistent results; however these studies have had small sample sizes, been conducted in single studies and have not sought independent replication.^15-17^ To our knowledge, no previous genome-wide association study (GWAS) of GWG has been conducted.

The aim of this study was to increase understanding of the genetic and non-genetic determinants of GWG by (a) estimating the proportion of variation in total, early and late GWG tagged by maternal and fetal common genetic variants; (b) undertaking a GWAS of maternal and fetal genetic variants with total, early and late GWG, and attempting to replicate associations in independent samples, and (c) determining the associations of genetic variants from GWAS of phenotypes that are plausible contributors to GWG (i.e. birthweight, BMI, waist-hip ratio, height, blood pressure, glucose, type 2 diabetes and vitamin D) with total, early and late GWG. We examined associations of maternal and fetal genetic exposures with total, early and late GWG, because the relative contribution of maternal and fetal phenotypes to GWG vary across gestation. For example, maternal fat deposition contributes relatively more to early GWG (up to ~ 18-20 weeks of gestation), and fetal growth more to later GWG.^3,10^ We included vitamin D (25(OH)D) as a phenotype that plausibly contributes to GWG as maternal 25(OH)D may have a positive affect on birthweight,^13^ and therefore may have a positive association with GWG.

## Participants and Methods

We included singleton pregnancies of mother-offspring pairs of European origin from 20 pregnancy/birth cohorts, described in detail in **eSupplementary material** and **eTable 1**. Pregnancies that resulted in a miscarriage or stillbirth, those with a known congenital anomaly and those where delivery was preterm (before 37 completed weeks of gestation) were excluded.

**Table 1.** Estimates of proportion of maternal and fetal genetic contributions from common variants to gestational weight gain.

|  | GREML Results<sup>a</sup> |  | M-GCTA Results |  |  |
| --- | --- | --- | --- | --- | --- |
|  | Maternal genome<br>N=6,435 | Child genome<br>N=6,418 | Maternal genome<br>N=4,078 | Child genome<br>N=4,078 | Covariance<br>N=4,078 |
| Early | 0.195 (0.055) | 0.058 (0.053) | 0.021 (0.113) | 0.000 (0.115) | 0.067 (0.091) |
| Late | 0.244 (0.054) | 0.110 (0.053) | 0.196 (0.113) | 0.161 (0.114) | -0.039 (0.091) |
| Total | 0.239 (0.055) | 0.121 (0.053) | 0.173 (0.112) | 0.045 (0.113) | 0.016 (0.090) |
| Birthweight | 0.13 (0.06) | 0.18 (0.06) | 0.04 (0.10) | 0.24 (0.11) | 0.04 (0.08) |

<sup>a</sup> P-values for the GREML results are: Maternal genome, early GWG = $1.12 \times 10^{-4}$ , Maternal genome, late GWG = $8.83 \times 10^{-7}$ , Maternal genome, total GWG = $1.94 \times 10^{-6}$ , Maternal genome, birth weight = 0.02, Offspring genome, early GWG = 0.130, Offspring genome, late GWG = 0.015, Offspring genome, total GWG = 0.008, Offspring genome, birth weight = $1.86 \times 10^{-3}$

Total GWG was defined as the last gestational weight (as long as this was ≥ 28 weeks of gestation) before delivery minus pre-/early-pregnancy weight divided by the length of gestation in weeks at the last measurement. Pre/early-pregnancy weight was defined as maternal self-reported pre-pregnancy weight (with this report collected during pregnancy), a research/clinical measure of weight prior to pregnancy (with that measure taken no more than 12-weeks before predicted date of conception), or the first antenatal clinic weight (with that assessment ≤ 13 weeks of gestation), which ever was the earliest. Early GWG was the difference between pre-/early-pregnancy weight and weight measured any time between 18 and 20 (inclusive) completed weeks of gestation divided by length of gestation in weeks at the time of the 18 to 20 week measurement. Late GWG was the difference between the 18 to 20 week measurement and the last gestational weight measure at ≥ 28 weeks of gestation divided by the gestational age difference in completed weeks between these two measurements. The gestational ages used to define early and late GWG were based on evidence regarding the different contributions of maternal fat deposition and fetal growth, with the former contributing relatively more to GWG up to ~ 18-20 weeks of gestation,^3,10^ and the latter more so after that point. Furthermore, applying multilevel models to the very detailed repeat measurements of gestational weight in the Avon Longitudinal Study of Parents and Children (ALSPAC), in which the median (IQR) of measurements per woman was 12 (9 to 13), demonstrated changes in the amount of weight gained per week of gestation at 19 and 28 weeks.^5^ In studies with repeated measurements between 18 and 20 weeks the one nearest to 18 weeks was used and in those with repeated later measurements the last weight was defined as the one nearest to (but before) delivery. Total, early and late GWG standard deviation (z-) scores were calculated within each study as the participant value minus the individual study mean then divided by the study standard deviation.

### Proportion of variation in total, early and late GWG, and birthweight that is due to maternal and fetal common genetic variants

We used methods that have been developed for use with genome-wide data to estimate the proportion of variation in total, early and late GWG, and birthweight tagged by maternal and fetal common genetic variants. Genetic restricted maximum likelihood (GREML)^18^ and maternal- genome-wide complex trait analysis (M-GCTA)^19^ were applied to maternal and fetal genome-wide data from ALSPAC. ALSPAC is a prospective population-based birth cohort study that recruited 14,541 pregnant women resident in Avon, UK with expected dates of delivery between 1^st^ April 1991 and 31^st^ December 1992 (http://www.alspac.bris.ac.uk.).^20,21^ GWG was determined using data extracted from obstetric medical records by trained research midwives.^5^ Birthweight, gestational age (in completed weeks) and fetal sex were obtained from obstetric/perinatal records. Maternal genome-wide data were obtained from the genome-wide Illumina 610 Quad Array. Fetal genome-wide data were obtained from the genome-wide Illumina 550 Quad Array. Further details, including genotype imputation and QC are provided in online **eSupplementary material** and characteristics of the participants are described in **eTable 1**.

### Maternal and fetal GWAS of total, early and late GWG

All studies in the Early Growth Genetics (EGG) consortium (http://egg-consortium.org/) with relevant data participated. Twenty independent pregnancy/birth cohorts contributed to at least one discovery and/or replication analysis (this included data from ALSPAC). Details of each of these studies are provided in **eSupplementary Material** and study participant characteristics, including their contribution to each GWAS, are shown in **eTable 1**. For total GWG, up to 10,543 and 16,317 participants contributed to maternal and fetal discovery GWAS, respectively, with numbers for early and late GWG GWAS being somewhat lower (**Table 2**). Up to an additional 10,660 and 7,561 participants contributed to maternal and fetal replication samples, with the maximum total meta-analysis sample size for total GWG being 18,420 and 21,105 for maternal and fetal GWAS, respectively.

**Table 2.**
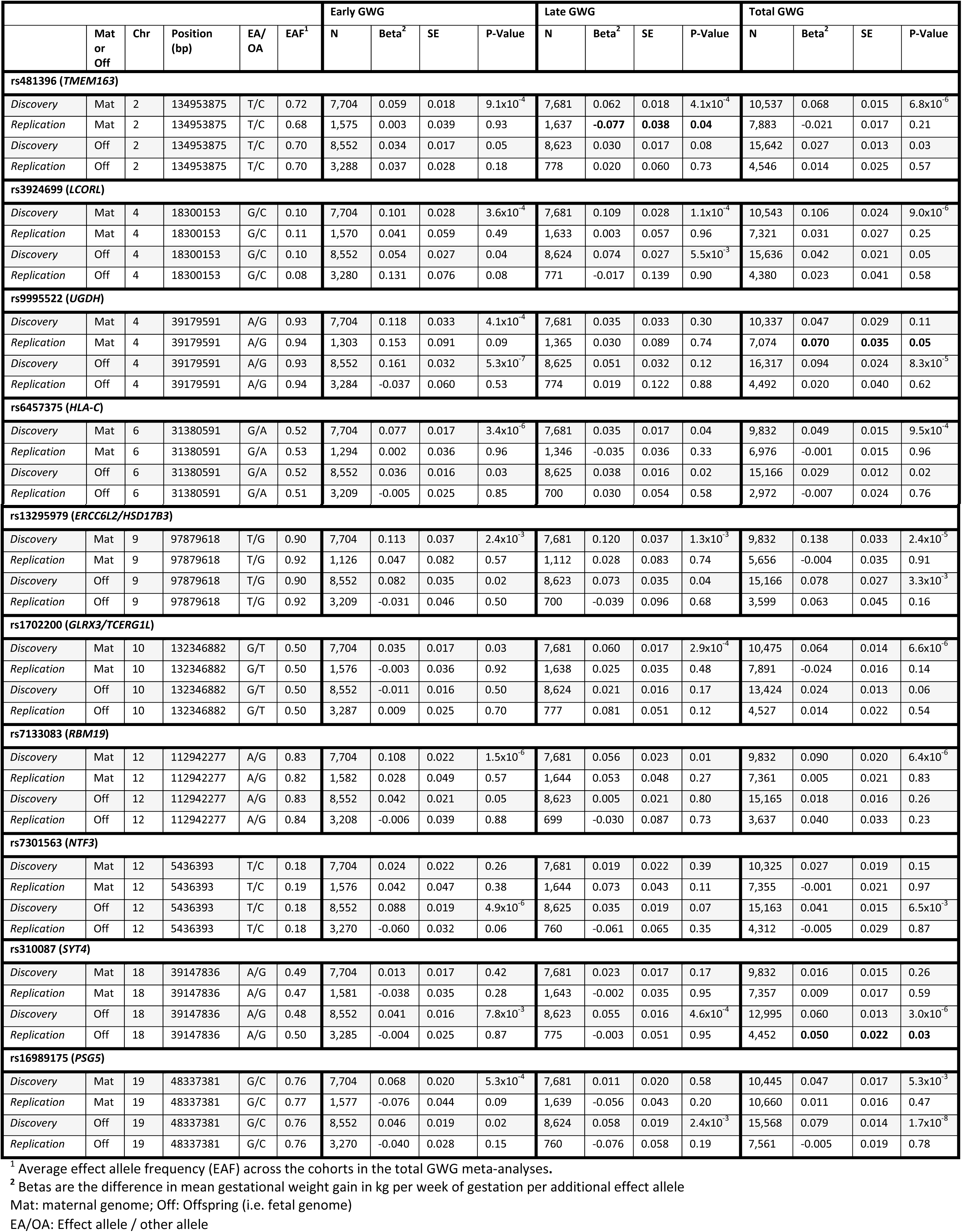
The most significant SNPs from each locus that reached P<10^−5^ from the discovery meta-analysis in all individuals. The nearest gene is used as the locus name.

GWAS discovery and replication analyses were undertaken independently by analysts working with each of the contributing studies following a prior agreed analysis plan. Genotypic data imputed to HapMap Phase 2 (Build 36, release 22) was used (methods for imputing within each contributing study is described in the **eSupplementary Material**) in the analysis, assuming an additive genetic model and adjusting for fetal sex.

Fixed-effects, inverse-variance weighted (IVW) meta-analyses in METAL^22^ were undertaken to combine GWAS results from the individual discovery studies. The most significant SNPs in regions reaching suggestive significance (P<10^−5^) in the discovery GWAS of any of the analyses (i.e. total, early or late GWG or in the maternal or fetal genome) were taken forward to replication. This set of SNPs were analysed against the three phenotypes in the replication studies and the results were combined using IVW meta-analysis in R (version 3.0.0) using the rmeta package.^23^ Additionally, to investigate whether this set of top SNPs were more likely to be acting in the maternal or offspring genome to influence GWG, conditional analysis was conducted in studies where both maternal and fetal genotype were available. Again, the results from these analyses were combined using IVW meta-analysis in R (version 3.0.0).

### Maternal and fetal genetic variants for phenotypes with plausible contributions to GWG

We examined the associations of a set of *a priori* agreed genetic variants that had previously (in GWAS) been shown to be robustly associated with phenotypes that might plausibly influence GWG with our GWG phenotypes. These were genetic variants for birthweight, BMI, waist-hip ratio, height, blood pressure, fasting glucose, type 2 diabetes and vitamin D (25(OH)D). **eTable 2** lists the variants included for each of the traits. The results for each of the variants were extracted from the discovery GWAS, and the replication studies provided results for the subset of variants they had available. IVW meta-analysis was conducted in METAL ^22^ to combine the results across all the cohorts. In these hypothesis driven analyses we use a two-sided p-value of < 0.05 as indicating statistical significance.

## Results

Total GWG was between 0.35 and 0.45kg/week in all of the general population studies and somewhat lower in the two studies that combined severely obese women with lean or a population cohort comparison group; in all studies early GWG was considerably lower than late GWG (**eTable 1**).

### Proportion of variation in GWG and birthweight due to maternal and fetal common genetic variants

SNPs across the genome explained broadly similar proportions of variation in late GWG and early GWG, but with stronger contributions of maternal compared with fetal genome, with SNPs in the maternal genome explaining approximately twice the amount of variation in total GWG than the fetal genome (**Table 1**). The opposite pattern was seen for birthweight, for example, SNPs across the maternal genome explained 24% (P = 1.94x10^−6^) of the variation in total GWG, with 12% (P = 0.008) explained by SNPs in the fetal genome, whereas the maternal genome explained 13% (P=0.02) and fetal genome 18% (P= 1.86×10^−3^) of variation in birthweight (**Table 1**). When we modelled maternal and fetal contributions together this pattern remained, but with the differences between maternal and fetal contributions increasing somewhat; for total GWG 17% and 5%, respectively for maternal and fetal genome and for birthweight 4% and 24%, respectively for maternal and fetal genome, with relatively little covariance between the two genomes for either trait. When the covariance and offspring/maternal variance components were constrained to zero in the M-GCTA model, similar results to the GREML analysis were obtained (**eTable 3**).

**Table 3.** Results from the unconditional analysis and analysis conditional on offspring (row 2 of each SNP) or maternal genotype (row 4 of each SNP) for the most significant SNPs from each locus that reached P<10^−5^ from the discovery meta-analysis; results from the maternal and offspring genotypes are presented. The nearest gene is used as the locus name.

|  |  |  |  |  | Early GWG |  |  |  | Late GWG |  |  |  | Total GWG<sup>1</sup> |  |  |  |
| --- | --- | --- | --- | --- | --- | --- | --- | --- | --- | --- | --- | --- | --- | --- | --- | --- |
|  | Genome | Chr | Position (bp) | EA/OA | N | Beta<sup>1</sup> | SE | P-Value | N | Beta<sup>1</sup> | SE | P-Value | N | Beta | SE | P-Value |
| **rs481396 (TMEM163)** |  |  |  |  |  |  |  |  |  |  |  |  |  |  |  |  |
| Unconditional | Mat | 2 | 134953875 | T/C | 6,635 | 0.040 | 0.019 | 0.03 | 6,674 | 0.057 | 0.019 | 2.3x10<sup>-3</sup> | 12,844 | 0.036 | 0.013 | 7.0x10<sup>-3</sup> |
| Conditional | Mat | 2 | 134953875 | T/C | 6,103 | 0.041 | 0.022 | 0.06 | 6,165 | 0.050 | 0.022 | 0.03 | 10,079 | 0.031 | 0.017 | 0.07 |
| Unconditional | Off | 2 | 134953875 | T/C | 6,291 | 0.040 | 0.019 | 0.04 | 6,356 | 0.031 | 0.019 | 0.11 | 11,340 | 0.033 | 0.015 | 0.03 |
| Conditional | Off | 2 | 134953875 | T/C | 6,103 | 0.012 | 0.022 | 0.59 | 6,165 | 0.011 | 0.023 | 0.64 | 10,079 | 0.019 | 0.018 | 0.28 |
| **rs3924699 (LCORL)** |  |  |  |  |  |  |  |  |  |  |  |  |  |  |  |  |
| Unconditional | Mat | 4 | 18300153 | G/C | 6,630 | 0.068 | 0.030 | 0.02 | 6,670 | 0.075 | 0.030 | 0.01 | 12,830 | 0.055 | 0.021 | 7.6x10<sup>-3</sup> |
| Conditional | Mat | 4 | 18300153 | G/C | 6,091 | 0.052 | 0.035 | 0.14 | 6,155 | 0.050 | 0.035 | 0.15 | 10,031 | 0.030 | 0.027 | 0.28 |
| Unconditional | Off | 4 | 18300153 | G/C | 6,283 | 0.050 | 0.030 | 0.09 | 6,350 | 0.074 | 0.030 | 0.01 | 11,241 | 0.047 | 0.023 | 0.05 |
| Conditional | Off | 4 | 18300153 | G/C | 6,091 | 0.025 | 0.035 | 0.47 | 6,155 | 0.045 | 0.035 | 0.20 | 10,031 | 0.027 | 0.028 | 0.33 |
| **rs9995522 (UGDH)** |  |  |  |  |  |  |  |  |  |  |  |  |  |  |  |  |
| Unconditional | Mat | 4 | 39179591 | A/G | 6,638 | 0.113 | 0.035 | 1.5x10<sup>-3</sup> | 6,677 | -0.020 | 0.026 | 0.44 | 12,838 | 0.055 | 0.026 | 0.03 |
| Conditional | Mat | 4 | 39179591 | A/G | 6,101 | 0.041 | 0.042 | 0.34 | 6,163 | 0.039 | 0.043 | 0.36 | 10,057 | 0.037 | 0.033 | 0.26 |
| Unconditional | Off | 4 | 39179591 | A/G | 6,288 | 0.179 | 0.035 | 4.8x10<sup>-7</sup> | 6,354 | -0.007 | 0.036 | 0.84 | 11,289 | 0.070 | 0.026 | 6.4x10<sup>-3</sup> |
| Conditional | Off | 4 | 39179591 | A/G | 6,101 | 0.163 | 0.041 | 8.2x10<sup>-5</sup> | 6,163 | -0.024 | 0.042 | 0.56 | 10,057 | 0.060 | 0.031 | 0.05 |
| **rs6457375 (HLA-C)** |  |  |  |  |  |  |  |  |  |  |  |  |  |  |  |  |
| Unconditional | Mat | 6 | 31380591 | G/A | 5,474 | 0.057 | 0.019 | 3.1x10<sup>-3</sup> | 5,446 | 0.009 | 0.020 | 0.64 | 9,530 | 0.020 | 0.014 | 0.16 |
| Conditional | Mat | 6 | 31380591 | G/A | 5,378 | 0.052 | 0.022 | 0.02 | 5,366 | 0.014 | 0.022 | 0.52 | 8,688 | 0.029 | 0.017 | 0.09 |
| Unconditional | Off | 6 | 31380591 | G/A | 5,480 | 0.023 | 0.020 | 0.233 | 5,458 | 0.020 | 0.007 | 0.74 | 9,388 | 0.005 | 0.015 | 0.71 |
| Conditional | Off | 6 | 31380591 | G/A | 5,378 | -0.0005 | 0.022 | 0.98 | 5,366 | -0.002 | 0.023 | 0.93 | 8,688 | -0.011 | 0.017 | 0.53 |
|  | Genome | Chr | Position (bp) | EA/OA | N | Beta<sup>1</sup> | SE | P-Value | N | Beta<sup>1</sup> | SE | P-Value | N | Beta | SE | P-Value |
| rs13295979 (ERCC6L2/HSD17B3) |  |  |  |  |  |  |  |  |  |  |  |  |  |  |  |  |
| Unconditional | Mat | 9 | 97879618 | T/G | 5,826 | 0.117 | 0.040 | 3.7x10<sup>-3</sup> | 5,805 | 0.118 | 0.041 | 3.9x10<sup>-3</sup> | 11,087 | 0.061 | 0.027 | 0.03 |
| Conditional | Mat | 9 | 97879618 | T/G | 5,577 | 0.086 | 0.046 | 0.06 | 5,564 | 0.105 | 0.047 | 0.02 | 9,367 | 0.055 | 0.034 | 0.10 |
| Unconditional | Off | 9 | 97879618 | T/G | 5,668 | 0.095 | 0.041 | 0.02 | 5,643 | 0.103 | 0.042 | 0.01 | 10,051 | 0.094 | 0.030 | 2.1x10<sup>-3</sup> |
| Conditional | Off | 9 | 97879618 | T/G | 5,577 | 0.051 | 0.046 | 0.26 | 5,564 | 0.052 | 0.046 | 0.27 | 9,367 | 0.061 | 0.035 | 0.08 |
| rs1702200 (GLRX3/TCERG1L) |  |  |  |  |  |  |  |  |  |  |  |  |  |  |  |  |
| Unconditional | Mat | 10 | 132346882 | G/T | 6,636 | 0.008 | 0.017 | 0.65 | 6,675 | 0.044 | 0.017 | 0.01 | 12,840 | 0.027 | 0.012 | 0.03 |
| Conditional | Mat | 10 | 132346882 | G/T | 6,105 | 0.003 | 0.021 | 0.90 | 6,167 | 0.054 | 0.021 | 0.01 | 10,082 | 0.032 | 0.016 | 0.05 |
| Unconditional | Off | 10 | 132346882 | G/T | 6,290 | -0.011 | 0.018 | 0.54 | 6,356 | 0.022 | 0.018 | 0.23 | 11,328 | 0.021 | 0.014 | 0.13 |
| Conditional | Off | 10 | 132346882 | G/T | 6,105 | 0.001 | 0.021 | 0.97 | 6,167 | -0.019 | 0.021 | 0.37 | 10,082 | -0.001 | 0.016 | 0.97 |
| rs7133083 (RBM19) |  |  |  |  |  |  |  |  |  |  |  |  |  |  |  |  |
| Unconditional | Mat | 12 | 112942277 | A/G | 5,834 | 0.099 | 0.025 | 5.3x10<sup>-5</sup> | 5,812 | 0.042 | 0.025 | 0.09 | 11,147 | 0.040 | 0.017 | 0.02 |
| Conditional | Mat | 12 | 112942277 | A/G | 5,580 | 0.105 | 0.029 | 2.7x10<sup>-4</sup> | 5,566 | 0.036 | 0.029 | 0.21 | 9,430 | 0.042 | 0.022 | 0.06 |
| Unconditional | Off | 12 | 112942277 | A/G | 5,667 | 0.033 | 0.025 | 0.20 | 5,642 | 0.026 | 0.025 | 0.31 | 10,089 | 0.038 | 0.019 | 0.05 |
| Conditional | Off | 12 | 112942277 | A/G | 5,580 | -0.011 | 0.029 | 0.70 | 5,566 | 0.001 | 0.029 | 0.96 | 9,430 | 0.022 | 0.022 | 0.32 |
| rs7301563 (NTF3) |  |  |  |  |  |  |  |  |  |  |  |  |  |  |  |  |
| Unconditional | Mat | 12 | 5436393 | T/C | 6,636 | 0.041 | 0.022 | 0.07 | 6,681 | 0.023 | 0.022 | 0.30 | 12,841 | 0.021 | 0.016 | 0.20 |
| Conditional | Mat | 12 | 5436393 | T/C | 6,084 | -0.012 | 0.026 | 0.64 | 6,146 | 0.013 | 0.027 | 0.63 | 9,965 | 0.002 | 0.021 | 0.94 |
| Unconditional | Off | 12 | 5436393 | T/C | 6,274 | 0.105 | 0.022 | 3.2x10<sup>-6</sup> | 6,340 | 0.028 | 0.023 | 0.22 | 11,100 | 0.064 | 0.018 | 3.3x10<sup>-4</sup> |
| Conditional | Off | 12 | 5436393 | T/C | 6,084 | 0.089 | 0.026 | 7.5x10<sup>-4</sup> | 6,146 | 0.044 | 0.027 | 0.10 | 9,965 | 0.071 | 0.021 | 8.7x10<sup>-4</sup> |
|  | Genome | Chr | Position (bp) | EA/OA | N | Beta<sup>1</sup> | SE | P-Value | N | Beta<sup>1</sup> | SE | P-Value | N | Beta | SE | P-Value |
| **rs310087 (SYT4)** |  |  |  |  |  |  |  |  |  |  |  |  |  |  |  |  |
| *Unconditional* | Mat | 18 | 39147836 | A/G | 6,641 | 0.012 | 0.017 | 0.49 | 6,680 | 0.020 | 0.017 | 0.24 | 12,848 | 0.016 | 0.012 | 0.18 |
| *Conditional* | Mat | 18 | 39147836 | A/G | 6,104 | -0.009 | 0.020 | 0.66 | 6,166 | -0.001 | 0.021 | 0.95 | 10,059 | -0.014 | 0.016 | 0.39 |
| *Unconditional* | Off | 18 | 39147836 | A/G | 6,288 | 0.035 | 0.017 | 0.04 | 6,353 | 0.057 | 0.017 | 1.1x10<sup>-3</sup> | 11,278 | 0.058 | 0.014 | 1.6x10<sup>-5</sup> |
| *Conditional* | Off | 18 | 39147836 | A/G | 6,104 | 0.041 | 0.020 | 0.04 | 6,166 | 0.057 | 0.021 | 5.3x10<sup>-3</sup> | 10,059 | 0.067 | 0.016 | 3.1x10<sup>-5</sup> |
| **rs16989175 (PSG5)** |  |  |  |  |  |  |  |  |  |  |  |  |  |  |  |  |
| *Unconditional* | Mat | 19 | 48337381 | G/C | 6,637 | 0.039 | 0.020 | 0.06 | 6,676 | 0.003 | 0.020 | 0.87 | 16,162 | 0.029 | 0.013 | 0.02 |
| *Conditional* | Mat | 19 | 48337381 | G/C | 6,085 | 0.026 | 0.034 | 0.28 | 6,147 | -0.024 | 0.024 | 0.32 | 13,143 | 0.004 | 0.017 | 0.82 |
| *Unconditional* | Off | 19 | 48337381 | G/C | 6,273 | 0.057 | 0.021 | 7.4x10<sup>-3</sup> | 6,339 | 0.049 | 0.021 | 0.02 | 14,363 | 0.058 | 0.014 | 3.7x10<sup>-5</sup> |
| *Conditional* | Off | 19 | 48337381 | G/C | 6,085 | 0.042 | 0.024 | 0.08 | 6,147 | 0.050 | 0.025 | 0.04 | 13,143 | 0.055 | 0.017 | 9.6x10<sup>-4</sup> |

<sup>1</sup> Betas are the difference in mean gestational weight gain in kg per week of gestation per additional effect allele
Mat: maternal genome; Off: Offspring (i.e. fetal genome)
EA/OA: Effect allele / other allele

### Maternal and fetal GWAS of total, early and late GWG

There was no systematic inflation of the test statistics in the meta-analysis of approximately 2.5 million SNPs (λ_mat-early_=1.01, λ_mat-late_ = 1.01, λ_mat-total_ = 1.02, λ_off-early_ = 1.02, λ_off-late_ = 1.00, λ_off-total_ = 1.00; **eFigure 1**). In discovery analyses, one variant, rs16989175 near the pregnancy specific beta 1-glycoprotein 5 (*PSG5*) gene, reached conventional GWAS significance (<5x10^−8^) for fetal genetic association with total GWG; it also showed some evidence of fetal association with late (p = 2.4 × 10^−3^) and early (p = 0.02) GWG. However, it did not replicate in either the maternal or fetal analysis. An additional 9 regions were identified as being suggestively significant (p < 10^−5^) for at least one phenotype in either maternal or fetal genome (**eFigure 2**). These were taken forward in replication analyses. Only one of these 10 SNPs replicated, rs310087 near *SYT4* (**Table 2**). This SNP was associated with total GWG in the fetal genome (mean difference in total GWG per allele 0.06 (95%CI: 0.04, 0.08) kg/week; p = 3×10^−6^ in discovery samples and 0.05 (95%CI: 0.01, 0.09) kg/week; p = 0.03 in replication samples and 0.06 (95%CI: 0.04, 0.08) kg/week; p = 1.6×10^−5^ in pooled discovery and replication). For six out of the ten top SNPs identified (rs481396, rs3924699, rs6457375, rs13295979, rs1702200 and rs7133083), the point estimate was larger for the maternal genotype on total GWG than the offspring genotype on this phenotype (**Table 2**). This was suggested in conditional analyses, whereby the point estimates for the offspring genotypes mostly attenuated after adjusting for maternal genotype (**Table 3**). The variant near *SYT4* that in fetal genome was nominally significantly associated with total GWG in discovery analyses and replicated, was not notably altered with adjustment for the maternal variant.

### Association of maternal and fetal genotypes for phenotypes with plausible contributions with GWG

Seven of the 32 BMI associated SNPs showed evidence of association (p < 0.05) with early GWG using maternal genotype, with five of the BMI increasing alleles associated with a decrease in early GWG (**eFigure 3**). In contrast, only three of the BMI associated SNPs showed association with late GWG using maternal genotype, and all three increased both BMI and GWG. A similar pattern of association was seen with the SNPs associated with glucose (**eFigure 4**) and type 2 diabetes (**eFigure 5**), whereby alleles associated with increased glucose/risk of type 2 diabetes showed evidence of association of maternal genotype with decreased GWG in early gestation and with increased GWG in late gestation. A smaller portion of SNPs for these phenotypes using the offspring genotype were associated with GWG (**eFigures 3 to 5**).

Surprisingly, none of the birthweight associated SNPs using the offspring genotype were associated with any GWG phenotype (**eFigure 6**). However, the SNP with the largest effect on birthweight (from <u>fetal</u> GWAS of birthweight), rs900400, using the maternal genotype was associated with decreased late GWG and total GWG, for each birthweight increasing SNP. SNPs associated with blood pressure, when using the maternal genome, showed stronger association with late GWG than early GWG, with the blood pressure increasing allele for most variants associating with decreased GWG (**eFigure 7**). Offspring blood pressure SNPs and SNPs associated with waist-hip ratio (maternal or offspring; **eFigure 8**), vitamin D (25(OH)D) (**eFigure 9**) or height (**eFigure 10**) were not notably associated with any GWG phenotypes.

## Discussion

We have shown that approximately 20% of the variability in GWG can be explained by common maternal genetic variants. A much smaller contribution is made by common genetic variants from the fetal genome. This pattern of maternal and fetal genetic contribution is opposite to what we see with birthweight, for which the fetal contribution is greater. Despite this modest genetic contribution, which is similar to the common genetic contribution to birthweight and many other phenotypes,^24^ in what we believe to be the first genome-wide association study of GWG, we were unable to identify any genetic variants that reached genome-wide levels of significance (P<5x10^−8^) and that replicated. Given the possible contribution of several adiposity related phenotypes to overall GWG, we also investigated whether genetic variants that are known to be associated with these traits were also associated with GWG. Some maternal BMI, fasting glucose and type 2 diabetes variants were nominally associated with GWG, such that those that were associated with increased BMI, glucose or type 2 diabetes, were associated with lower early and higher late GWG. Some maternal variants associated with higher systolic blood pressure also associated with lower late GWG. In general fetal variants associated with these traits were largely unrelated to GWG. Of note, established maternal and fetal birthweight variants were for the most part not related to GWG. The one exception being rs900400, a variant previously shown to be strongly related to birthweight in a genome-wide study of fetal genotype,^25^ which in our study was inversely associated with late and total GWG in the case of the maternal genotype. This variant has also been recently shown to be inversely associated with leptin in genome-wide analyses,^26^ and thus the inverse association of this variant in the mother with GWG may reflect a positive association of maternal leptin with GWG.

Using a twin study, Andersson et al show that the heritability of first pregnancy GWG is 43%;^14^ we were able to show that approximately half of this could be explained by common genetic variants or variants they tag in the maternal genome. This is similar to the proportion of heritability explained in other common traits such as height and BMI.^24^ It is perhaps not surprising that our results suggest that the maternal genome has a greater contribution to GWG than the offspring genome. On average, approximately 55% of GWG is a result of increased maternal tissue, 15-20% is due to the placenta and amniotic fluid, and 20-25% is a result of fetal tissue.^27^ The maternal genome will contribute to tissue expansion in the mother, as well as to placental size, amniotic fluid and fetal growth, whereas, it is likely that the fetal genome will only contribute to placenta, amniotic fluid and fetal growth. We detected some evidence of a negative genetic covariance in the M-GCTA analysis of late GWG. A negative covariance implies that a proportion of maternal genetic variants associated with increased GWG are associated with decreased GWG when present in the fetal genome. Although this was a surprising result, it is not inconceivable. For example, there is a well described relationship between mutations in the glucokinase gene (GCK) and offspring birthweight, whereby if the mutation is present in the mother and not the offspring then birthweight is increased, whereas if the mutation is present in the offspring but not the mother then birthweight is decreased.^28^ Given birthweight is a component of GWG, it is plausible that variants in *GCK* and other mutations involved in insulin secretion could produce similar effects on GWG. However, given the large standard error on the estimate, this negative covariance might be a chance finding and requires replication before any further interpretation is made.

Our lack of replicated genome-wide significant findings might be due to the complexity of the GWG phenotype. Weight was measured by trained personnel during pregnancy in the majority of studies included in this meta-analysis, however the pre-pregnancy measure was often self-report and the early pregnancy measure would have included some pregnancy weight gain. This would have increased the measurement error for GWG, making it difficult to identify true genetic associations. In addition, we had low statistical power to detect associations with genetic variants which have a small effect. With an alpha of 5x10^−8^ in the maternal GWAS of total GWG, we had 80% power to detect a genetic variant that explained between 0.37% and 0.4% of the variance for our range of sample sizes (N=9,832 – 10,543). Similarly, we had 80% power to detect a variant that explained 0.24% - 0.3% of the variance in the offspring GWAS of total GWG (N=12,995 – 16,317). However, for other complex quantitative phenotypes, such as BMI, the genetic variants discovered to date each explain 0.003-0.325% of the variance,^29^ indicating that many common genetic variants each of small effect influence the trait. Therefore, we had adequate power to detect common genetic variants with modest to large effects, but we were unable to detect variants with smaller effects, even though we used the largest sample of individuals for exploring genetic associations with this phenotype to date and are unaware currently of other European origin studies that could have added to this effort.

Despite most of our analyses suggesting a stronger contribution of maternal, than fetal common genetic variants to GWG, the one nominally significant variant that replicated was for a fetal variant that was related to total GWG. This variant on chromosome 18, is near to the Synaptotagmin 4 (*SYT4*) gene, which is a protein coding gene involved in calcium and syntaxin binding.^30^ Its relation to GWG is unclear and this association should be treated with caution unless further replicated, particularly as the association was only nominal and did not reach conventional GWAS significance of 5×10^−8^ even in combined meta-analysis of discovery and replication samples (the result being per allele difference in mean total GWG: 0.06 (95%CI: 0.04, 0.08) kg/week; p = 1.6 ×10^−5^ in pooled discovery and replication).

We expanded on previous studies that examined the associations of candidate maternal and/or fetal adiposity or diabetes related genetic variants with GWG^15-17^ by looking at a wider variety of phenotypes that are observationally related to GWG (previous studies looked only at BMI and type 2 diabetes), in a considerably larger sample of participants and using a greater number of variants for each phenotype. We are aware that for some of the phenotypes investigated, there are a larger number of associated SNPs in the more recent GWAS, for example over 90 variants have now been shown to independently relate to BMI.^29^ The subset of variants that we used for each phenotype were those with the largest effect sizes on each of the individual traits and that were available in the majority of replication (as well as discovery) samples, therefore we will have greater power to detect an effect with GWG if one exists.

The main strength of this study was the availability of both maternal and offspring genotype and the three separate phenotypes for GWG allowing us to investigate whether genetic variants had consistent effects throughout pregnancy. The choice of 18-20 weeks to distinguish between early and late GWG was determined through multilevel modelling in the ALSPAC cohort, which was the largest contributing cohort to our study with multiple repeated weight measures throughout pregnancy, and availability of data from the other cohorts involved.^5^ Despite being the first large GWAS of this trait to our knowledge and our effort to include all studies of European origin women with relevant data we had limited power to detect variants with weak effects and will continue to seek additional studies to contribute to large GWAS in the future.

In summary, we have identified that a substantial proportion of the variation in GWG can be explained by common variants in the maternal genome, with an additional smaller proportion being explained by the offspring genome. In what we believe to be the first GWAS of GWG, using the largest collection of individuals, we were unable to identify any loci with a large effect on GWG, but found some further evidence that maternal variants may contribute more to GWG than fetal variants. These initial results suggest that the association of GWG with later offspring outcome may reflect intrauterine (maternal) effects. However, given the composite nature of GWG, including increasing maternal fat stores and plasma volume, the growing fetus, placenta and amniotic fluid, larger sample sizes are required to identify individual genetic loci for GWG.

## References

1. Rasmussen KM, Yaktine AL. Weight gain during pregnancy: reexamining the guidelines. Committee to reexamine IOM pregnancy weight guidelines: US Institute of Medicine and National Research Council; 2009.

2. Viswanthan M, Siega-Riz AM, Moos M-K, et al. Outcomes of Maternal Weight Gain, Evidence Report/Technology Assessment No. 168. AHRQ Publication No08-E009 edRockville, MD: Agency for Healthcare Research and Quality 2008.

3. Lawlor DA, Relton C, Sattar N, Nelson SM. Maternal adiposity--a determinant of perinatal and offspring outcomes? Nat Rev Endocrinol 2012; 8(11): 679-88.

4. Macdonald-Wallis C, Tilling K, Fraser A, Nelson SM, Lawlor DA. Gestational weight gain as a risk factor for hypertensive disorders of pregnancy. Am J Obstet Gynecol 2013; 209(4): 327 e1-17.

5. Fraser A, Tilling K, Macdonald-wallis C, et al. Association of maternal weight gain in pregnancy with offspring obesity and metabolic and vascular traits in childhood. Circulation 2010; 121: 2557-64.

6. Lawlor DA, Lichtenstein P, Fraser A, Langstrom N. Does maternal weight gain in pregnancy have long-term effects on offspring adiposity? A sibling study in a prospective cohort of 146,894 men from 136,050 families. AM J CLIN NUTR 2011; 94(1): 142-8.

7. Mamun AA, O’Callaghan M, Callaway L, Williams G, Najman J, Lawlor DA. Associations of gestational weight gain with offspring body mass index and blood pressure at 21 years of age: evidence from a birth cohort study. Circulation 2009; 119(13): 1720-7.

8. Fraser A, Tilling K, Macdonald-Wallis C, et al. Associations of gestational weight gain with maternal body mass index, waist circumference, and blood pressure measured 16 y after pregnancy: the Avon Longitudinal Study of Parents and Children (ALSPAC). Am J Clin Nutr 2011; 93(6): 1285-92.

9. Lewis AJ, Galbally M, Gannon T, Symeonides C. Early life programming as a target for prevention of child and adolescent mental disorders. BMC Med 2014; 12: 33.

10. Lawlor DA. The Society for Social Medicine John Pemberton Lecture 2011. Developmental overnutrition--an old hypothesis with new importance? Int J Epidemiol 2013; 42(1): 7-29.

11. Davey Smith G, Hemani G. Mendelian randomization: genetic anchors for causal inference in epidemiological studies. Hum Mol Genet 2014; 23(R1): R89-R98.

12. Lawlor DA, Harbord RM, Sterne JAC, Timpson NJ, Davey Smith G. Mendelian randomization: using genes as instruments for making causal inferences in epidemiology. Statistic in Medicine 2008; 27: 1133-63.

13. Tyrrell J, Richmond RC, Palmer TM, et al. Genetic Evidence for Causal Relationships Between Maternal Obesity-Related Traits and Birth Weight. JAMA 2016; 315(11): 1129-40.

14. Andersson ES, Silventoinen K, Tynelius P, Nohr EA, Sorensen TI, Rasmussen F. Heritability of gestational weight gain--a Swedish register-based twin study. Twin research and human genetics: the official journal of the International Society for Twin Studies 2015; 18(4): 410-8.

15. Lawlor DA, Fraser A, Macdonald-Wallis C, et al. Maternal and offspring adiposity-related genetic variants and gestational weight gain. Am J Clin Nutr 2011; 94(1): 149-55.

16. Stuebe AM, Lyon H, Herring AH, et al. Obesity and diabetes genetic variants associated with gestational weight gain. Am J Obstet Gynecol 2010; 203(3): 283-17.

17. Dishy V, Gupta S, Landau R, et al. G-protein beta(3) subunit 825 C/T polymorphism is associated with weight gain during pregnancy. Pharmacogenetics 2003; 13(4): 241-2.

18. Yang J, Benyamin B, McEvoy BP, et al. Common SNPs explain a large proportion of the heritability for human height. Nat Genet 2010; 42(7): 565-9.

19. Eaves LJ, Pourcain BS, Davey Smith G, York TP, Evans DM. Resolving the effects of maternal and offspring genotype on dyadic outcomes in genome wide complex trait analysis (“M-GCTA”). Behav Genet 2014; 44(5): 445-55.

20. Boyd A, Golding J, Macleod J, et al. Cohort Profile: The ‘Children of the 90s’--the index offspring of the Avon Longitudinal Study of Parents and Children. Int J Epidemiol 2013; 42(1): 111-27.

21. Fraser A, Macdonald-Wallis C, Tilling K, et al. Cohort Profile: The Avon Longitudinal Study of Parents and Children: ALSPAC mothers cohort. Int J Epidemiol 2013; 42(1): 97-110.

22. Willer CJ, Li Y, Abecasis GR. METAL: fast and efficient meta-analysis of genomewide association scans. Bioinformatics 2010; 26(17): 2190-1.

23. Ihaka R, Gentleman R. R: a language for data analysis and graphics. Journal of Computational and Graphical Statistics 1996; 5: 299-314.

24. Yang J, Manolio TA, Pasquale LR, et al. Genome partitioning of genetic variation for complex traits using common SNPs. Nat Genet 2011; 43(6): 519-25.

25. Horikoshi M, Yaghootkar H, Mook-Kanamori DO, et al. New loci associated with birth weight identify genetic links between intrauterine growth and adult height and metabolism. Nat Genet 2013; 45(1): 76-82.

26. Kilpelainen TO, Carli JF, Skowronski AA, et al. Genome-wide meta-analysis uncovers novel loci influencing circulating leptin levels. Nature communications 2016; 7: 10494.

27. Pitkin RM. Nutritional support in obstetrics and gynecology. ClinObstetGynecol 1976; 19(3): 489-513.

28. Hattersley AT, Beards F, Ballantyne E, Appleton M, Harvey R, Ellard S. Mutations in the glucokinase gene of the fetus result in reduced birth weight. Nature Genetics 1998; 19(3): 268-70.

29. Locke AE, Kahali B, Berndt SI, et al. Genetic studies of body mass index yield new insights for obesity biology. Nature 2015; 518(7538): 197-206.

30. Adolfsen B, Littleton JT. Genetic and molecular analysis of the synaptotagmin family. Cellular and molecular life sciences: CMLS 2001; 58(3): 393-402.

